# Genetic Risk Factors for Colorectal Cancer in Multiethnic Indonesians

**DOI:** 10.1101/626739

**Authors:** Irawan Yusuf, Upik A. Miskad, Ronald E. Lusikooy, Arham Arsyad, Akram Irwan, George Mathew, Ivet Suriapranata, Rinaldy Kusuma, Bens Pardamean, Muhamad Fitra Kacamarga, Arif Budiarto, Tjeng Wawan Cenggoro, Carissa I. Pardamean, Christopher McMahan, Chase Joyner, James W. Baurley

## Abstract

**Purpose:** Colorectal cancer is a common cancer in Indonesia, yet it has been understudied. We conduct a genome-wide association study focused on evaluation and discovery of colorectal cancer risk factors in Indonesians.

**Methods:** We administered detailed questionnaires and collecting blood samples from 162 colorectal cancer cases throughout Makassar, Indonesia. We also established a control set of 193 healthy individuals frequency matched by age, sex, and ethnicity. A genome-wide association analysis was performed on 84 cases and 89 controls passing quality control. We evaluated known colorectal cancer genetic variants using logistic regression and established a genome-wide polygenic risk model using a Bayesian variable selection technique.

**Results:** We replicate associations for rs9497673, rs6936461 and rs7758229 on chromosome 6; rs11255841 on chromosome 10; and rs4779584, rs11632715, and rs73376930 on chromosome 15. Polygenic modeling identified 10 SNP associated with colorectal cancer risk.

**Conclusions:** This work helps characterize the relationship between variants in the *SCL22A3*, *SCG5*, *GREM1*, and *STXBP5-AS1* genes and colorectal cancer in a diverse Indonesian population. With further biobanking and international research collaborations, variants specific to colorectal cancer risk in Indonesians will be identified.

## 1 Introduction

Colorectal cancer is one of the most common cancers in the world and a leading cause of cancer-related deaths [1][2]. There is growing evidence that colorectal cancer rates are changing in Asian countries, but the causes are still under investigation [3][4]. Colorectal cancer is now one of the top three cancers in many Asian countries [5]. Currently, Asia contributes to 48% of the total number of new colorectal cancer cases in the world, of which the majority are found in Eastern Asia [6]. Specifically in Indonesia, the age-standardized incidence for males and females has been reported as 15.9 and 10.1 per 100,000 respectively [7].

The contribution of heritable factors towards colorectal cancer occurrence is estimated to be between 12-35%. However, germline mutations that are highly penetrant contribute less than 5% to colorectal cancer [8]. Nonetheless, increasing evidence is finding that heritability plays a potential, crucial role in colorectal cancer pathogenesis. Currently, mutations in 14 genes are suspected to underlie different subtypes of colorectal cancer, including mutations in the APC that increases predisposition to familial adenomatous polyposis (FAP) and defects in mismatch repair genes associated with Lynch Syndrome [8]. Recent genome-wide association studies have identified common genetic variants linked to colorectal cancer predisposition, highlighting a greater association between heritable risk and the disease. Thus far, over 40 genetic variants have been identified, within several well-known biological pathways that have been shown to be highly relevant to oncogenesis, including the TGF-beta/BMP pathway and the mitogen-activated protein kinases (MAPK) pathway [8].

However, many of these colorectal cancer genetic associations were discovered in European-ancestry populations but do not replicate well in other ancestry groups, demonstrating the need for studies in diverse populations worldwide [9]. The Asia Colorectal Cancer Consortium was initiated in 2009 among East Asian nations and has successfully identified novel relevant, genetic regions [10, 11]. However, colorectal cancer cases from South East Asian cohorts, have been under represented.

Given the changes in colorectal cancer rates in Asia and the differences in risk factors present in ethnically diverse South East Asia, we present results of the first genomic association study of colorectal cancer in Indonesia. We present results from the initial phase of this study, focused on cases from South Sulawesi, Indonesia.

## 2 Methodology

### 2.1 Study participants

162 colorectal cancer cases were recruited from 7 hospitals throughout Makassar, Indonesia between 2014 and 2016. The hospitals were Wahidin Sudirohusodo Hospital, Hasanuddin University Hospital, Ibnu Sina Hospital, Akademis Hospital, Grestelina Hospital, Stella Maris Hospital, and Hikmah Hospital. 193 controls were frequency matched to cases on age category, sex, and ethnicity. This research was approved by the Hasanuddin University Ethical Committee (registration number: UH 15040389).

### 2.2 Data and DNA sample collection

Questionnaires and medical records were recorded into study data collection forms and entered into a study database. The case forms contained 382 questions and the control forms contained 319 questions. The forms included information on demographics, cancer history in the family, smoking behavior, alcohol use, and detailed dietary history. For colorectal cancer cases, the forms collected information on cancer symptoms, staging (post operation), tumor, location, histopathology, and type of surgery. The database was managed by the Bioinformatics and Data Science Research Center (BDSRC) at Bina Nusantara University (Jakarta, Indonesia). A blood sample was collected from the basilic/cephalic vein on all participants for genotyping. These blood samples were stored in Hasanuddin University Laboratory at minus 20 degrees Celsius.

### 2.3 Genotyping and imputation

DNA was extracted from samples at Mochtar Riady Institute for Nanotechnology (MRIN) Laboratory (Tangerang, Indonesia). Genomic DNA was extracted from 200 µL of whole blood sample using the QIAamp DNA Mini Kit (Qiagen, Hilden, Germany) according to the manufacturer’s protocol. DNA concentration was determined using NanoDrop ND-1000 spectrophotometer, version 3.3 (Thermo Fisher Scientific, Wilmington, DE, USA) and adjusted to a concentration of 20 ng/µL. The quality of DNA extracted was verified by purity index of OD260/OD280 (1.8-2.0) and OD260/OD230 (> 1.5). The DNA was inspected through Gel Electrophoresis using 1% molecular biology grade Agarose (Biorad, Hercules, CA, USA). Extracted DNA were sent to RUCDR Infinite Biologics for genotyping (Piscataway, NJ, USA) under Material Transfer Agreement (MTA) approved by the Indonesian Health Ministry (registration number: LB.02.01/I/12749/2016).

DNA samples from study cases and controls were genome-wide genotyped on the Smokescreen Genotyping Array [12]. Using 200 ng of genomic DNA, array plates were prepared using the Axiom 2.0 Reagent Kits and then processed on the GeneTitan MC instrument (Thermo Fisher Scientific, Wilmington, DE, USA). Analysis of the raw data was performed using Affymetrix Power tools (APT) v-1.16 according to the Affymetrix best practices workflow. 183 samples remained after completing these steps. Additional steps were performed using SNPolisher to identify and select best performing probe sets and high quality SNPs for downstream analysis. 524,765 SNPs remained after QC filtering. Additional sample quality control included verifying concordance of study replicates, checking for unintentional duplicates and unexpected relatives, and verifying genetic versus reported gender. After filtering samples with missing covariates, 173 samples (84 cases and 89 controls) remained for statistical analysis.

Genome-wide imputation was performed on the Michigan Imputation Server v1.0.2 [13]. Briefly, quality controlled study genotypes were reported on the forward strand and uploaded in vcf format. 1000 Genomes Phase 3 [14] was selected as a reference panel, phasing was performed using Eagle v2.3 [15], and allele frequencies were compared against the 1000 Genomes East Asian (EAS) populations. The server automatically excludes variants with alleles other than (A,C,T,G), variants with duplicate positions, indels, monomorphic sites, and allele mismatches with the reference panel.

### 2.4 Statistical analysis

#### 2.4.1 Ancestry analysis

Ancestry categories were estimated from 5,515 ancestry informative markers contained on the Smokescreen Genotyping Array using fastStructure 1.0 [16]. Combining study and reference data from the 1000 Genomes Project Phase 3, we estimated the ancestry proportions of East Asian (EAS), South Asian (SAS), European (EUR), and African (AFR).

#### 2.4.2 Genome-wide association analysis

We filtered out variants with poor imputation quality (< 0.3) and rare variants (minor allele < 1%). We then performed a marginal analysis of the remaining SNP genotype dosages fitting logistic regression models, with sex, age, body mass index, smoking status and estimated ancestries proportions (i.e., SAS,EUR,AFR) as covariates. The threshold for statistical significance in the discovery scan was set at the historical traditional genome-wide value of 5E-8.

We queried the scan results for markers previously reported to be associated with colorectal cancer. These variants were identified through previous genotyping in an independent sample of South Sulawesi colorectal cancer cases (R. Kusuma, I. Suriapranata, personal communication) and a recent catalog of colorectal cancer SNPs for a genome-wide association scan in Hispanics [17]. The source and annotation for these variants are provided in Supplementary Table 3. Variants with evidence of replication (p-value < 0.05) were flagged for further investigation.

We also developed a polygenic model considering the joint effect of multiple genetic variants on colorectal cancer. We selected the top 200 SNPs, based on Bayes factors [18], as candidate predictors in this joint model. Bayes factors were computed for the marginal versus the null models for each SNP while controlling for gender, age, BMI, and smoking status. To jointly model these variants, we use a Bayesian variable selection technique. In particular, we fit a logistic regression model utilizing shrinkage priors for each of the explanatory variables; i.e., the covariates listed above as well as the remaining candidate SNPs. In this analysis, the generalized double Pareto shrinkage prior of [19] was specified and the parameters of the joint model were estimated via a maximum a posteriori (MAP) estimator [19] which was obtained via an expectation-maximization (EM) algorithm [20]. The MAP estimator under these specifications simultaneously completes parameter estimation and variable selection by obtaining a sparse estimator [21]; i.e., some of the regression coefficients are estimated to be identically equal to zero thus removing the effect of the corresponding explanatory variable. The EM algorithm was developed following the techniques illustrated by [19, 22] and the regularization parameters were selected via the Bayesian information criterion [23]. All statistical analysis was performed in R [24].

## 3 Results

### 3.1 Characteristics of study sample

The characteristics of the colorectal cancer cases and controls are summarized in Table 1. The mean age of the colorectal cancer cases was 54 years. The majority of cases were male (57%). Among ethnicities, most cases were self-reported Bugis (44%) or Makassar ethnicity (27%). Controls appeared to be adequately frequency matched to cases by age, sex, and ethnicity (*p* > 0.05). Colorectal cancer cases had lower average body mass index (BMI) and were more likely to be smokers than controls (*p* < 0.01). Estimated genetically, the majority of both cases and controls were of East Asian ancestry. 87% of the cases had late stage cancer (III or IV) which unfortunately is consistent with recent reports in Indonesia [25]. As seen in other studies, the most common colorectal cancer site was rectum (43%) [26, 27].

**Table 1.**
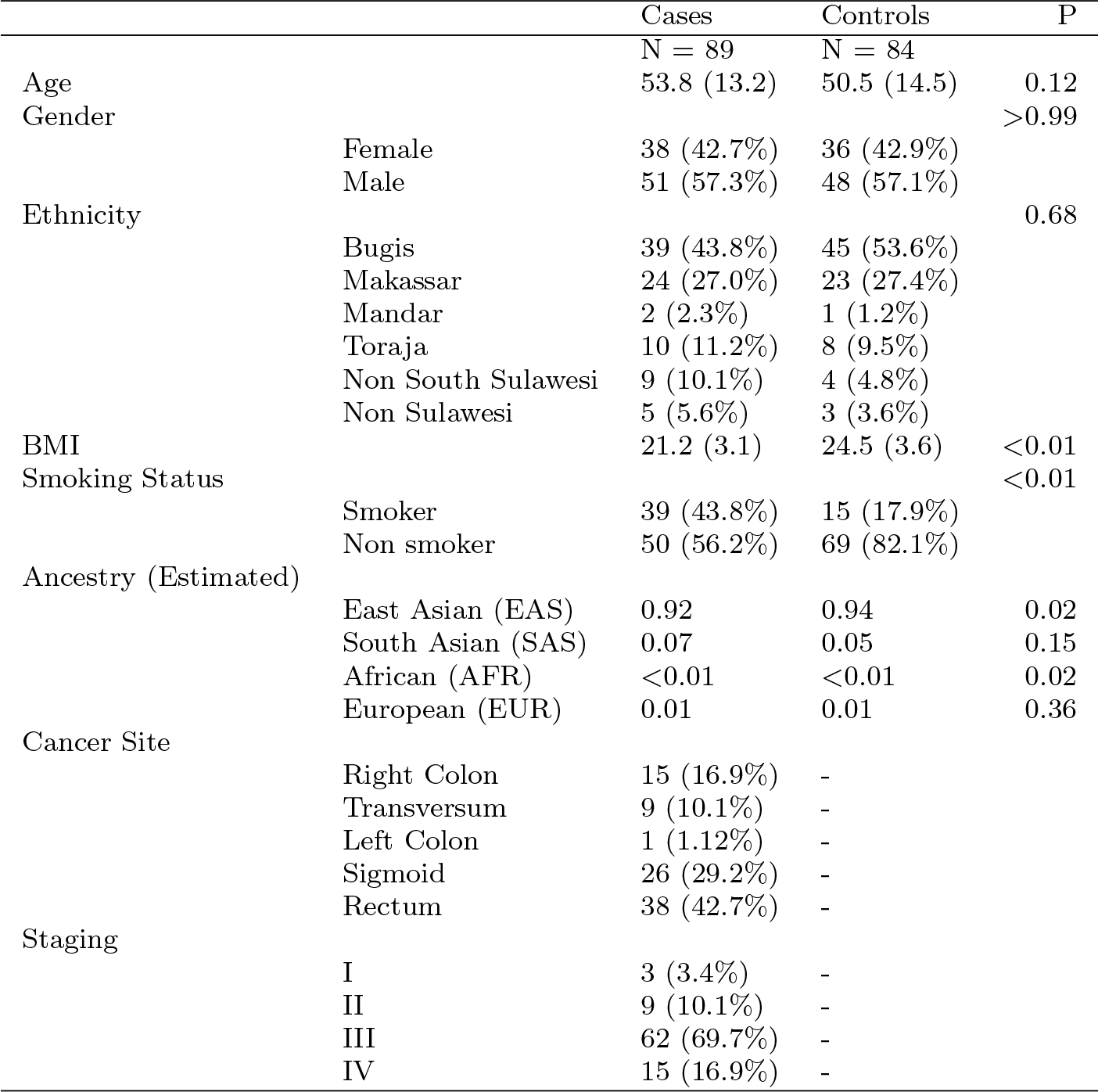
Characteristics of South Sulawesi colorectal cancer cases and controls

**Table 2.** Polygenic risk model learned from colorectal cancer data. Presented results include the chromosome (Chr) and position of the significant genetic variants, the gene they lie on (Gene), reference allele (Ref), minor allele frequency (MaF), and estimated effect (Estimate).

| Description | Chr | Position | Gene | Ref | MaF | Estimate |
| --- | --- | --- | --- | --- | --- | --- |
| Intercept |  |  |  |  |  | 0.90 |
| Gender |  |  |  |  |  | 0.00 |
| Age |  |  |  |  |  | -3.75 |
| BMI |  |  |  |  |  | 0.00 |
| Smoking |  |  |  |  |  | 1.32 |
| rs11919079 | 3 | 57086348 | Intron:ARHGEF3 | G | 0.07 | 2.40 |
| rs4888186 | 16 | 81947156 | Intron:PLCG2 | C | 0.08 | 0.85 |
| rs11016111 | 10 | 129963848 | Intergenic | C | 0.34 | -1.32 |
| rs77657157 | 5 | 98125016 | Intron:RGMB | G | 0.05 | 1.95 |
| - | 18 | 59822981 | Deletion:PIGN | TC | 0.19 | -1.39 |
| rs17066763 | 5 | 164113078 | Intergenic | T | 0.12 | 1.65 |
| rs2446103 | 6 | 77328692 | Intergenic | A | 0.04 | 1.22 |
| rs7219420 | 17 | 45800299 | Intergenic | T | 0.36 | 1.32 |
| - | 16 | 13018917 | Insertion:SHISA9 | C | 0.11 | 1.67 |
| rs78165118 | 3 | 12816282 | Intergenic | A | 0.03 | 2.13 |

### 3.2 Genome-wide association analysis

As expected given the sample size, no SNPs met the historical cutoff set for genome-wide significance (Supplementary Figures 6 and 7). The summaries for all variants with a marginal p-value < 5E 5 are included in the supplementary materials (Table 4). These include two intergenic SNPs and two SNPs in the *MRO* gene on chromosome 18.

Results for previously reported colorectal cancer SNPs are presented in Figure 1 and Supplementary Table 3. There is evidence of replication for the following genetic variants: rs9497673, rs6936461 and rs7758229 on chromosome 6; rs11255841 on chromosome 10; and rs4779584, rs11632715, and rs73376930 on chromosome 15. The regions are characterized in Figures 2, 3, 4, and 5. The pattern of associations is rather diffuse in the *STXBP5-AS1* (STXBP5 Antisense RNA 1) and *SLC22A3* genes of chromosome 6, representing the correlation among the variants in these regions (Figures 2 and 3). Similarly, the association pattern tapers along chromosome 10. The strongest association pattern can be found on chromosome 15. This region has a more defined peak than the other regions with associations spanning two genes: *SCG5* (secretogranin V) and *GREM1* (gremlin 1, DAN family BMP antagonist).

**Fig. 1.**
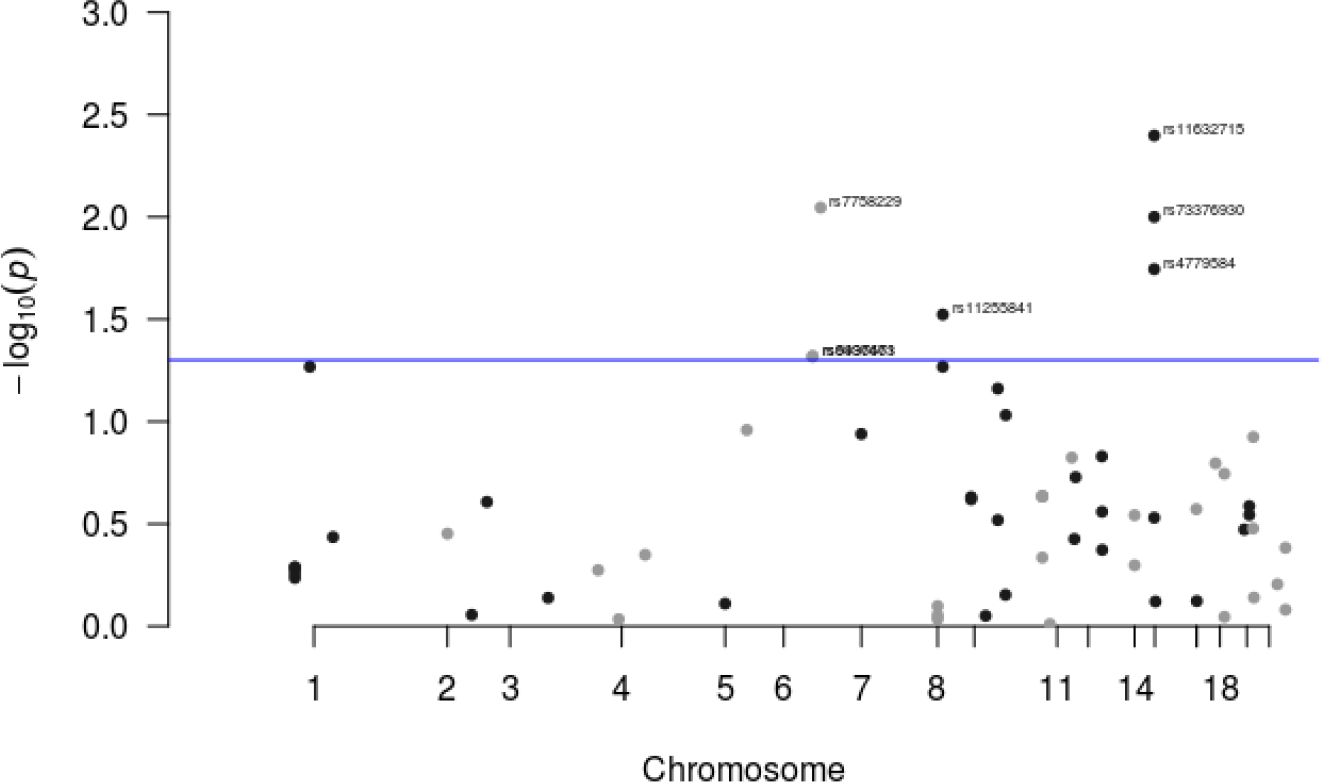
Results for known colorectal cancer susceptibility SNPs. Variants with p-values < 0.05 were flagged for further investigation.

**Fig. 2.**
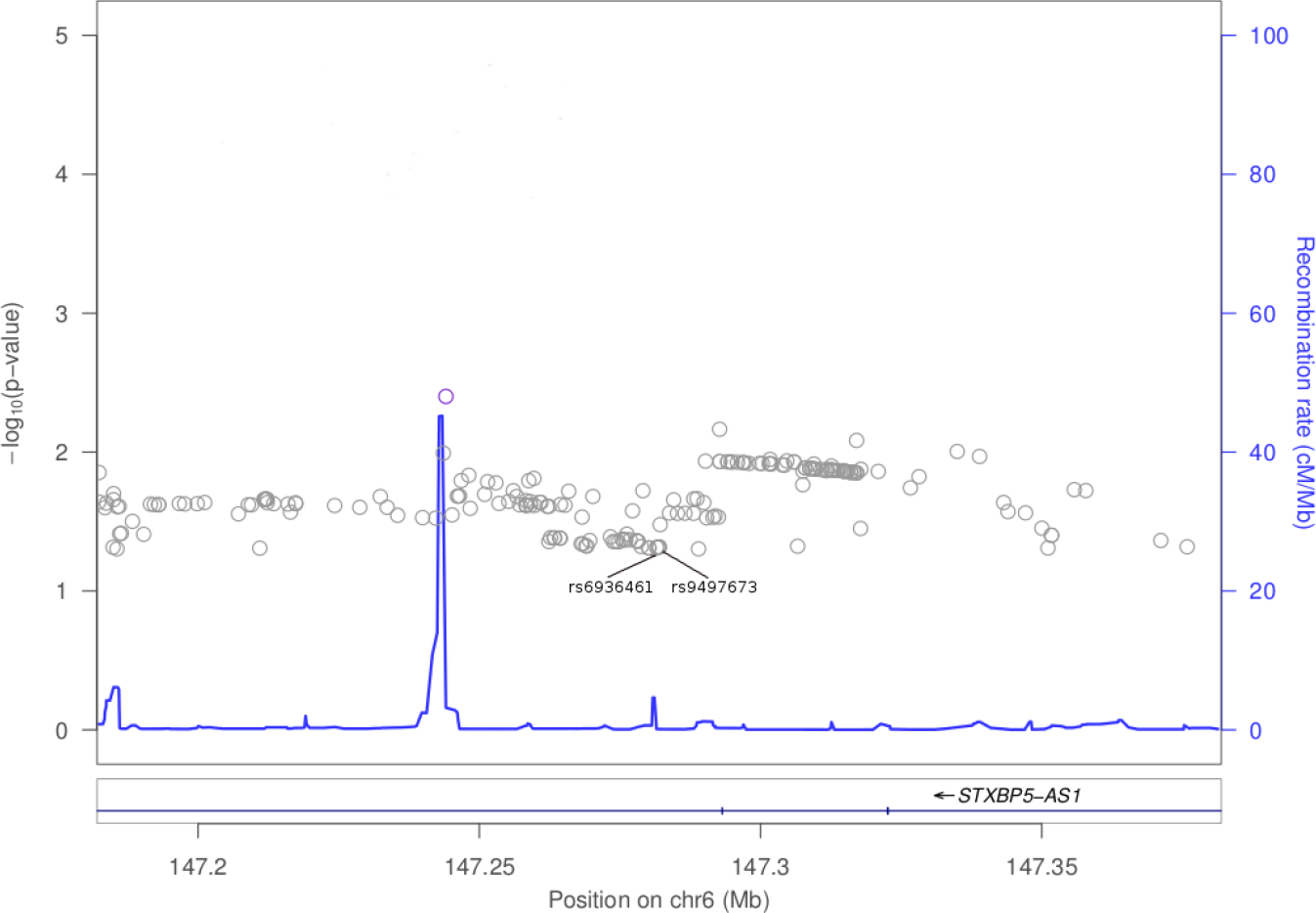
Association plot for 100kb region flanking rs6936461 on chromosome 6

**Fig. 3.**
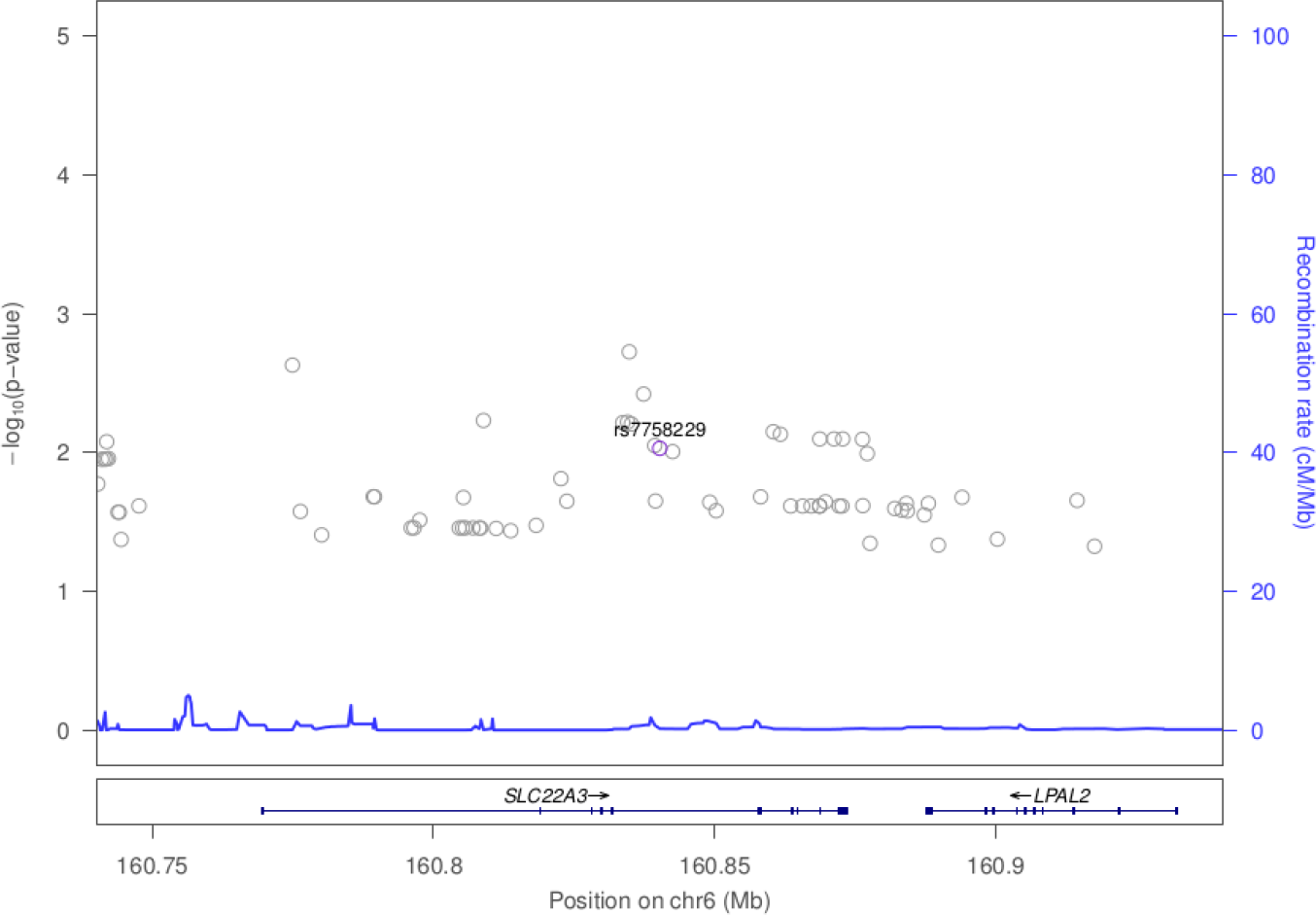
Association plot for 100kb region flanking rs7758229 on chromosome 6

**Fig. 4.**
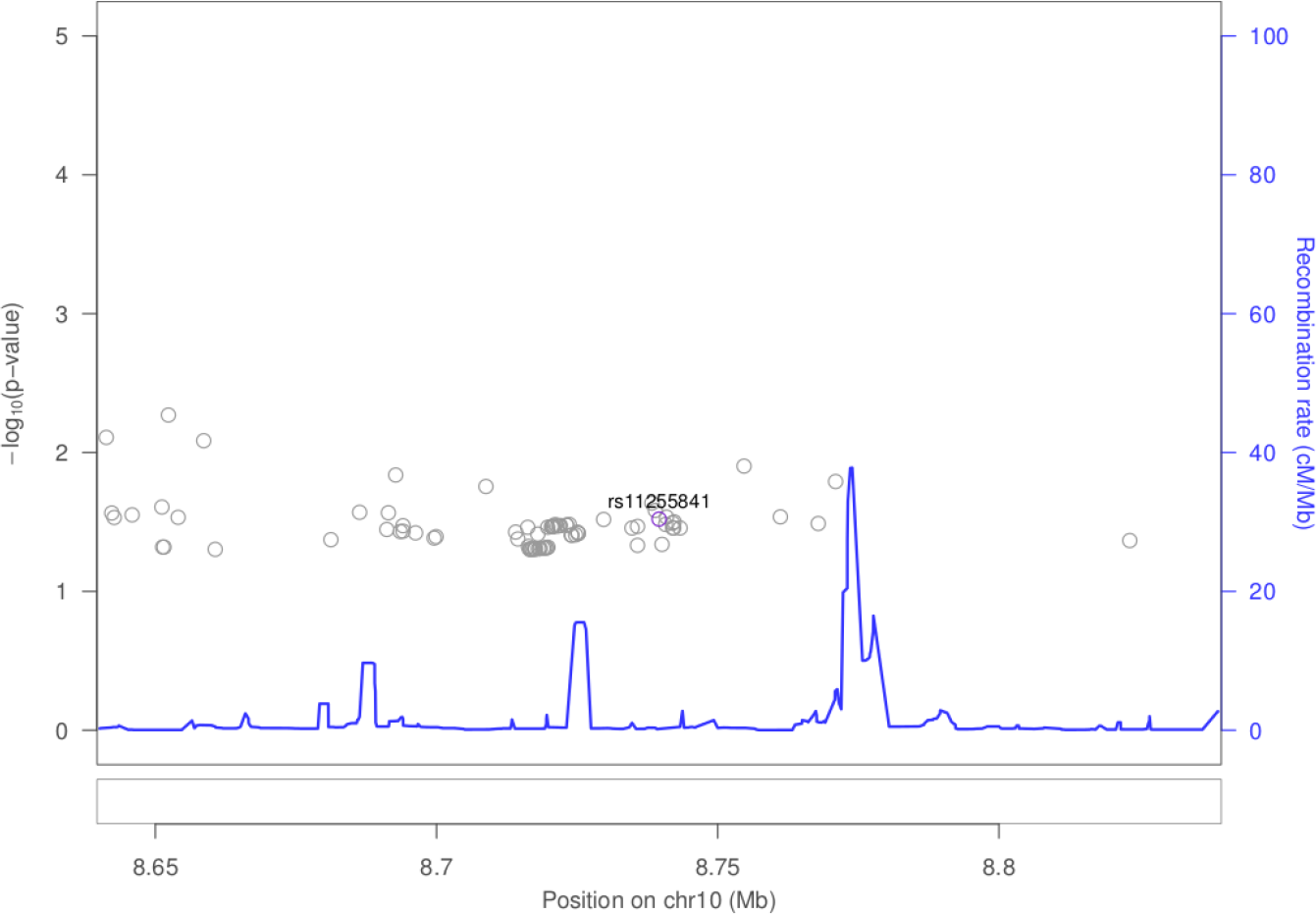
Association plot for 100kb region flanking rs11255841 on chromsome 10

**Fig. 5.**
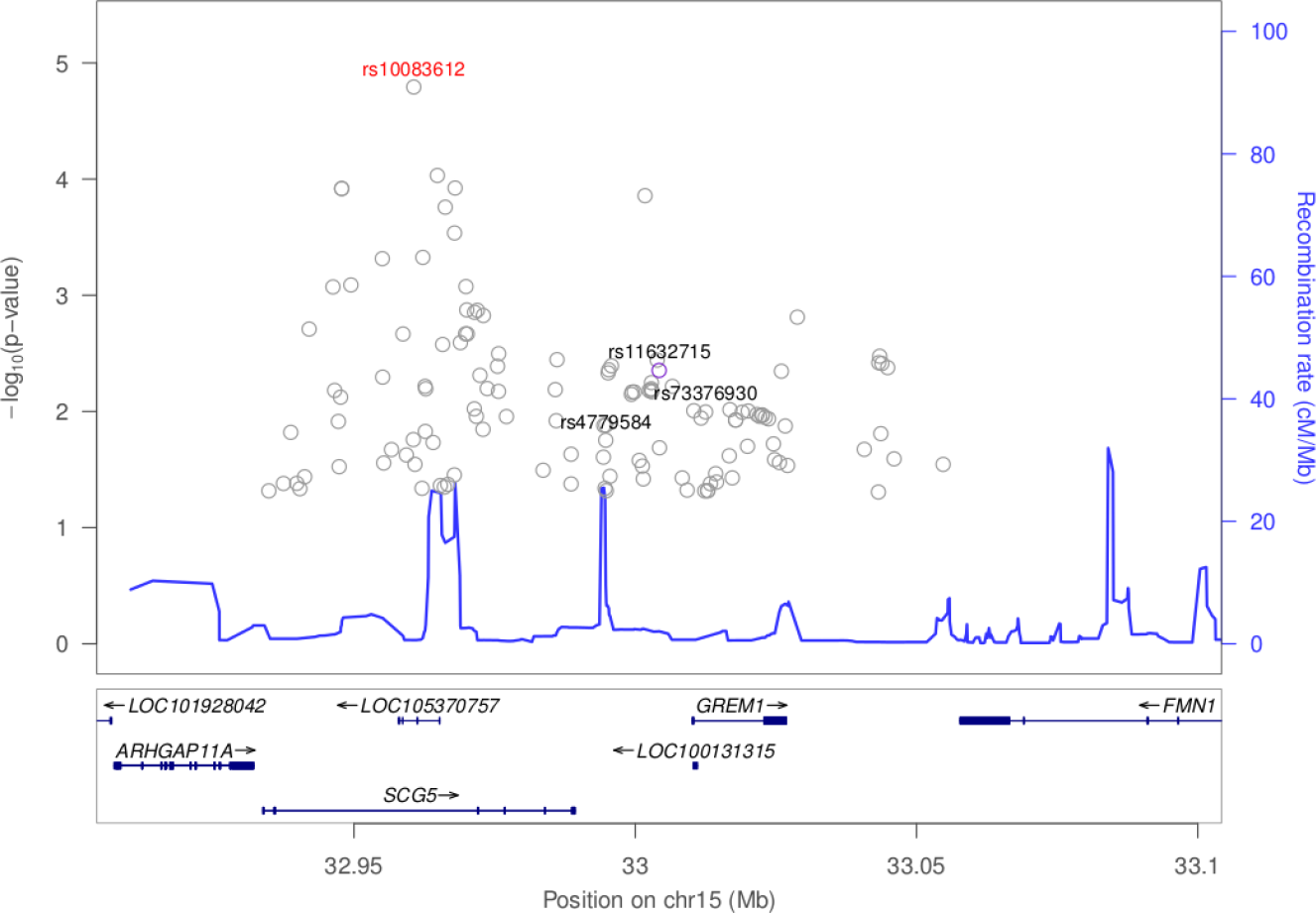
Association plot for 100kb flanking rs11632715 on chromosome 15. The top associated SNP in the region was rs10083612.

The polygenic analysis identified 10 SNPs which appear to have a relatively strong association (i.e., large effect size) with the risk of developing colorectal cancer. Five of these SNPs lie in intergenic regions; three lie in introns of *ARHGEF3*, *PLCG2*, and *RGMB*; one is a deletion in *PIGN*; and one is an insertion in *SHISA9*.

## 4 Discussion

This study represents the first genome-wide analysis of a South Sulawesi population in Indonesia. Strengths of the study include the building of a colorectal cancer research program in Indonesia, the extensive questionnaire for assessing non-genetic risk factors, and genome-wide genotyping across diverse ethnicities. Limitations of the study include the sample size, which restricts the analysis to previously identified colorectal cancer markers and challenges shared by case-control study designs. For instance, the controls may represent different groups than cases. We attempted to account for this by frequency matching on age, sex, and ethnicity. Additionally, the timing of assessments need to be considered in interpreting the results. Given screening programs are still being developed in Indonesia, the majority of the cases had late stage colorectal cancer, stage III and IV. When BMI was assessed in these patients they already had significant weight lose, thus the direction of the effect is different than what one might expect.

Several previously identified colorectal cancer associated SNPs replicated in this population. And we can begin characterizing these regions by examining neighboring variants.

rs7758229 within *SLC22A3* on chromosome 6 was originally identified and subsequently replicated in large case-control study of a Japanese population (OR of 1.3) [28]. Interestingly, in a subsequent study in a Chinese population, this SNP was not associated with colorectal cancer (OR of 0.95) [29]. However, in S. Sulawesi, we detect a statistically significant association with colorectal cancer (p=0.009, OR of 2.2). Given these difference among East Asians, further work to understand variation in *SLC22A3* and colorectal cancer is needed. *SLC22A3* encodes for the protein OCT3, which is an organic cationic transporter. While OCT3/SLC22A3 is well characterized within neurochemistry, it has been found to play a role within oncology as well. The upregulation of SLC22A3 in head and neck squamous cell carcinoma is associated with improved prognosis while the downregulation of SLC22A3 leads to enhanced metastasis and invasion of the tumor [30]. SLC22A3 has also been implicated in the pathogenesis of prostate cancer and its expression is elevated in these neoplastic tissues [31]. The level of OCT3/SLC22A3 expression has also been linked to the level of patient responsiveness towards cancer treatments [32]; in particular, platin-based cytotoxic cancer treatments in colorectal cancer [33] patients, as well as head and neck squamous cell carcinoma patients [30].

Intergenic variant rs11255841 on chromosome 10 was identified in an colorectal cancer GWAS of European ancestry individuals [34] and has replicated in a Japanese study and a large meta-analysis with nearly 37,000 cases [35, 36]. With the risk allele of T, this variant had an odds ratio of 2.2 in our study, while previous reports had an odds ratio of 1.1-1.2.

The region on chromosome 15 nearby *SCG5* and *GREM1* have been flagged in multiple GWAS, e.g., [37]. We replicated colorectal cancer associations for rs4779584 (p=0.018), rs11632715 (p=0.004), and rs73376930 (p=0.010). Interestingly, the smallest p-value in the region was rs10083612 within an intron of *SCG5* (p=1.61e-5, see Figure 5). The role of SCG5 in colorectal cancer has not been well characterized, while much is known about its neighbor *GREM1*’s role in colorectal cancer. GREM1, which is one of the antagonists of the bone morphogenetic proteins (BMPs) found within the TGF-beta signaling pathway, has been found to be important for the survival and proliferation of several types of cancers [38]. In particular, modulated expression of *GREM1* is found in cancer-associated stromal cells. GREM1 is also found to be a proangiogenic factor, suggesting a role in cancer development when it is upregulated [39]. SCG5 and GREM1 genes have been found to be associated with polyposis syndromes that are associated with colorectal cancer [40]. A duplication that spans the 3’end of *SCG5* and the immediate, adjacent upstream region of *GREM1* is associated with hereditary mixed polyposis syndrome (HMPS) as well as tumorigenesis in juvenile polyposis. This duplication results in a 40-kb extra segment that leads to the upregulation of *GREM1* expression. The duplication is the basis for an autosomal dominant HMPS condition that is prevalent among the Ashkenazi Jewish population and is a recommended biomarker/genetic test to detect CRC in this population. Aberrant expression of *GREM1* has also been shown to underlie oncogenesis within the large intestines and colon [41].

Two of the previously identified colorectal cancer markers replicate in this study (rs6936461 and rs9497673; see Table 3). These SNPs are located in the intronic regions of *STXBP5-AS1* on chromosome 6. Using bioinformatics tools, it is predicted that changes from T to A in rs6936461 and A to G in rs9497673, has the potential to alter the splicing of the gene [42]. *STXBP5-AS1* is an long non-coding (lncRNA) gene. lncRNAs drive many important cancer phenotypes through their interactions with other cellular macromolecules including DNA, protein, microRNA and mRNA. The different expression of lncRNAs in colorectal cancer indicate that lncRNAs are involved in all stages of colorectal cancer. In colorectal cancer pathogenesis, lncRNAs are implicated in a variety of signaling pathways including the Wnt/-catenin signaling pathway, epidermal growth factor receptor (EGFR)/insulin-like growth factor type I receptor (IGF-IR) signaling pathway, KRAS and phosphatidylinositol-3-kinase (PI3K) pathways, transforming growth factor-beta (TGF-) signaling pathway, p53 signaling pathway, and the epithelial-mesenchymal transition (EMT) pathway [43]. While it is still unclear how *STXBP5-AS1* contributes to colon carcinogenesis, in a study involving 1067 breast cancer samples, Guo et al. identified *STXBP5-AS1* among lncRNA genes which play a role in predicting the prognostic survival with good sensitivity and specificity. The lncRNAs may act as competing endogenous RNAs (ceRNAs) and interfere in the binding of miR-190b to certain targets such as ERG, STK38L, and FNDC3A and thus contribute to breast cancer pathogenesis [44]. *STXBP5-AS1* may act similarly in colorectal cancer; it may hinder the binding of microRNAs to their target genes and subsequently modulate colorectal cancer tumorigenesis.

Interestingly, STXBP5-AS1 was identified among genes that are methylated in buccal samples in a genome-wide screen for cigarette smoke exposure, indicating its possible role in smoking-related diseases [45]. Since there is a significant difference in smoking status between cases and controls in our cohort, it is plausible that genetic variants associated with tobacco smoke are also associated with the presence of colorectal cancer in our study population.

The polygenic model represents a strategy for jointly modeling SNP effects in a GWAS and development of risk prediction models in a specific population. These models can be used to estimate an individuals risk of colorectal cancer based on easily obtainable genotypes. While most of the variants flagged in the polygenic model are novel, the gene *ARHGEF3* has been implicated in promoting nasopharyngeal carcinoma in Asians [46]. *RGMB* has been shown to promote colorectal cancer growth [47]. Additional samples will enable us to refine and validate a polygenic colorectal cancer risk model in Indonesians.

## 5 Conclusion

We demonstrate replication of several colorectal cancer genetic risk factors in an Indonesian population. This study provides rational for additional data collection in this population to characterize these regions more precisely and identify genetic risk factors unique to this diverse population.

## Acknowledgements

We would like to acknowledge Bina Nusantara and Hasanuddin University for funding this study, MRIN Laboratory for DNA Extraction, RUCDR Infinite Biologics for DNA processing and genotyping, BioRealm for support of the Smokescreen Genotyping Array, Research credits from Amazon Web Services (AWS) and generous contributions from NVIDIA and the AI R&D Center at Bina Nusantara University for computing and database support.

## 8 Supplementary materials

**Table 3.** Results for previously identified colorectal cancer SNPs

| Rsid | Gene | Chr | Pos | Ref | Alt | Source | MaF | OR | SE | P |
| --- | --- | --- | --- | --- | --- | --- | --- | --- | --- | --- |
| rs12124798 | Intron:RP11-451O13.1 | 1 | 157906955 | A | T | [48] | 0.091 | 1.309 | 0.489 | 0.581 |
| rs6684686 | Intron:RP11-451O13.1 | 1 | 157909708 | A | G | [48] | 0.091 | 1.327 | 0.472 | 0.549 |
| rs6701170 | Intron:RP11-451O13.1 | 1 | 157913954 | T | C | [48] | 0.090 | 1.362 | 0.476 | 0.516 |
| rs4971169 | Intron:RP11-451O13.1 | 1 | 157914330 | T | C | [48] | 0.088 | 1.372 | 0.486 | 0.516 |
| rs10911251 | Intron:LAMC1 | 1 | 183081194 | A | C | [49, 50] | 0.345 | 0.503 | 0.356 | 0.054 |
| rs6687758 | Intergenic | 1 | 222164948 | A | G | [51] | 0.093 | 1.550 | 0.486 | 0.367 |
| rs11903757 | Intergenic | 2 | 192587204 | T | C | [49] | 0.161 | 0.713 | 0.364 | 0.353 |
| rs35360328 | Intergenic | 3 | 40924962 | T | A | [52] | 0.142 | 1.079 | 0.505 | 0.880 |
| rs812481 | Intron:LRIG1 | 3 | 66442435 | C | G | [52] | 0.158 | 0.607 | 0.432 | 0.247 |
| rs10936599 | Synonymous:MYNN | 3 | 169492101 | C | T | [51] | 0.437 | 1.104 | 0.284 | 0.727 |
| rs7356196 | Intergenic | 4 | 84173104 | G | A | [48] | 0.309 | 1.229 | 0.329 | 0.532 |
| rs3987 | Intron:AC108056.1 | 4 | 118759055 | A | G | [53] | 0.396 | 1.028 | 0.292 | 0.925 |
| rs35509282 | Intergenic | 4 | 163333405 | T | A | [17] | 0.320 | 0.784 | 0.321 | 0.448 |
| rs647161 | Intron:CTC-203F4.1—CTC-349C3.1 | 5 | 134499092 | C | A | [10] | 0.419 | 0.923 | 0.284 | 0.776 |
| rs1321311 | Intron:PI16 | 6 | 36622900 | C | A | [54] | 0.332 | 0.621 | 0.298 | 0.110 |
| rs9497673 | Intron:STXBP5-AS1 | 6 | 147281518 | G | A | [48] | 0.289 | 1.955 | 0.340 | 0.048 |
| rs7758229 | Intron:SLC22A3 | 6 | 160840252 | G | T | [55] | 0.428 | 2.244 | 0.311 | 0.009 |
| rs10499807 | Intergenic | 7 | 68459734 | C | T | [48] | 0.347 | 1.792 | 0.370 | 0.115 |
| rs10505477 | Intron:RP11-382A18.1 | 8 | 128407443 | A | G | [56, 57] | 0.301 | 0.954 | 0.325 | 0.884 |
| rs6983267 | Intron:RP11-382A18.1 | 8 | 128413305 | G | T | [56, 58, 59, 60, 55] | 0.303 | 0.922 | 0.318 | 0.798 |
| rs7014346 | Intron:RP11-382A18.1 | 8 | 128424792 | A | G | [61, 51] | 0.254 | 0.966 | 0.350 | 0.922 |
| rs10795668 | Intergenic | 10 | 8701219 | G | A | [59] | 0.249 | 0.525 | 0.334 | 0.054 |
| rs11255841 | Intergenic | 10 | 8739580 | T | A | [50] | 0.254 | 0.462 | 0.357 | 0.030 |
| rs10763129 | Intron:PCDH15 | 10 | 56468045 | T | G | [48] | 0.124 | 0.557 | 0.498 | 0.240 |
| rs10825383 | Intron:PCDH15 | 10 | 56473754 | G | A | [48] | 0.124 | 0.560 | 0.488 | 0.234 |
| rs704017 | Intron:RP11-202P11.1 | 10 | 80819132 | A | G | [11] | 0.272 | 0.958 | 0.311 | 0.890 |
| rs1035209 | Intergenic | 10 | 101345366 | C | T | [50] | 0.159 | 2.105 | 0.410 | 0.069 |
| rs11190164 | Intergenic | 10 | 101351704 | A | G | [52] | 0.251 | 1.460 | 0.367 | 0.303 |
| rs12241008 | Intron:VTI1A | 10 | 114280702 | T | C | [62] | 0.335 | 0.895 | 0.290 | 0.703 |
| rs11196172 | Intron:TCF7L2 | 10 | 114726843 | G | A | [11] | 0.481 | 1.674 | 0.307 | 0.093 |
| rs174537 | Intron:C11orf9—RP11-467L20.9 | 11 | 61552680 | G | T | [11] | 0.081 | 0.684 | 0.518 | 0.462 |
| rs4246215 | Utr3:FEN1 | 11 | 61564299 | G | T | [11] | 0.087 | 0.543 | 0.510 | 0.232 |
| rs174550 | Utr5:FADS1 | 11 | 61571478 | T | C | [11] | 0.087 | 0.543 | 0.510 | 0.232 |
| rs1535 | Intron:FADS2 | 11 | 61597972 | A | G | [11] | 0.087 | 0.543 | 0.510 | 0.232 |
| rs3824999 | Intron:POLD3 | 11 | 74345550 | T | G | [54] | 0.376 | 0.991 | 0.295 | 0.977 |
| rs3802842 | Intron:C11orf92—C11orf93 | 11 | 111171709 | C | A | [61] | 0.324 | 0.651 | 0.298 | 0.150 |
| rs10774214 | Intron:RP11-264F23.3—RP11-264F23.4 | 12 | 4368352 | T | C | [10] | 0.253 | 0.737 | 0.343 | 0.375 |
| rs10849432 | Intergenic | 12 | 6385727 | C | T | [11] | 0.130 | 0.562 | 0.437 | 0.187 |
| rs34245511 | Intron:LIMA1 | 12 | 50573433 | G | C | [50] | 0.191 | 0.600 | 0.353 | 0.148 |
| rs7136702 | Intergenic | 12 | 50880216 | T | C | [51] | 0.382 | 1.333 | 0.263 | 0.276 |
| rs11169552 | Intergenic | 12 | 51155663 | C | T | [51] | 0.474 | 1.243 | 0.273 | 0.424 |
| rs4444235 | Intergenic | 14 | 54410919 | T | C | [51, 59] | 0.497 | 1.352 | 0.283 | 0.287 |
| rs1957636 | Intergenic | 14 | 54560018 | T | C | [59] | 0.350 | 1.267 | 0.354 | 0.504 |
| rs16969681 | Intergenic | 15 | 32993111 | C | T | [59] | 0.445 | 1.356 | 0.291 | 0.295 |
| rs4779584 | Intergenic | 15 | 32994756 | T | C | [63, 59] | 0.130 | 0.358 | 0.433 | 0.018 |
| rs11632715 | Intergenic | 15 | 33004247 | G | A | [59] | 0.241 | 4.728 | 0.546 | 0.004 |
| rs73376930 | Intron:GREM1 | 15 | 33012502 | A | G | [50] | 0.446 | 3.010 | 0.428 | 0.010 |
| rs1851317 | Intron:RP11-814P5.1 | 15 | 35077786 | A | C | [48] | 0.423 | 1.100 | 0.309 | 0.758 |
| rs9929218 | Intron:CDH1 | 16 | 68820946 | G | A | [51] | 0.410 | 0.729 | 0.285 | 0.268 |
| rs12603526 | Intron:NXN | 17 | 800593 | T | C | [11] | 0.133 | 0.882 | 0.399 | 0.754 |
| rs12458173 | Intergenic | 18 | 31430167 | G | A | [48] | 0.309 | 1.845 | 0.436 | 0.160 |
| rs7229639 | Intron:SMAD7 | 18 | 46450976 | A | G | [11] | 0.124 | 1.079 | 0.612 | 0.901 |
| rs4939827 | Intron:SMAD7 | 18 | 46453463 | T | C | [64, 61] | 0.312 | 0.645 | 0.327 | 0.180 |
| rs10411210 | Intron:RHPN2 | 19 | 33532300 | C | T | [51] | 0.153 | 0.705 | 0.365 | 0.337 |
| rs1800469 | Intron:TMEM91 | 19 | 41860296 | A | G | [11] | 0.498 | 1.359 | 0.288 | 0.286 |
| rs2241714 | Nonsynonymous:B9D2 | 19 | 41869392 | T | C | [11] | 0.491 | 1.364 | 0.275 | 0.259 |
| rs961253 | Intergenic | 20 | 6404281 | C | A | [51, 59] | 0.179 | 0.693 | 0.379 | 0.333 |
| rs4813802 | Intergenic | 20 | 6699595 | T | G | [59, 49] | 0.239 | 0.594 | 0.334 | 0.119 |
| rs2423279 | Intergenic | 20 | 7812350 | T | C | [10] | 0.416 | 1.106 | 0.287 | 0.725 |
| rs6066825 | Intron:PREX1 | 20 | 47340117 | A | G | [52] | 0.416 | 0.844 | 0.346 | 0.624 |
| rs4925386 | Intron:LAMA5 | 20 | 60921044 | T | C | [51, 49] | 0.260 | 0.758 | 0.338 | 0.414 |
| rs2427308 | Intron:CABLES2 | 20 | 60969451 | C | T | [50] | 0.190 | 1.141 | 0.619 | 0.831 |

Chr: Chromosome
Pos: Chromosome Position (build 37)
Ref/Alt: Reference and alternate allele
MaF: Minor allele frequency
OR: Odds ratio
SE: Standard error

**Fig. 6.**
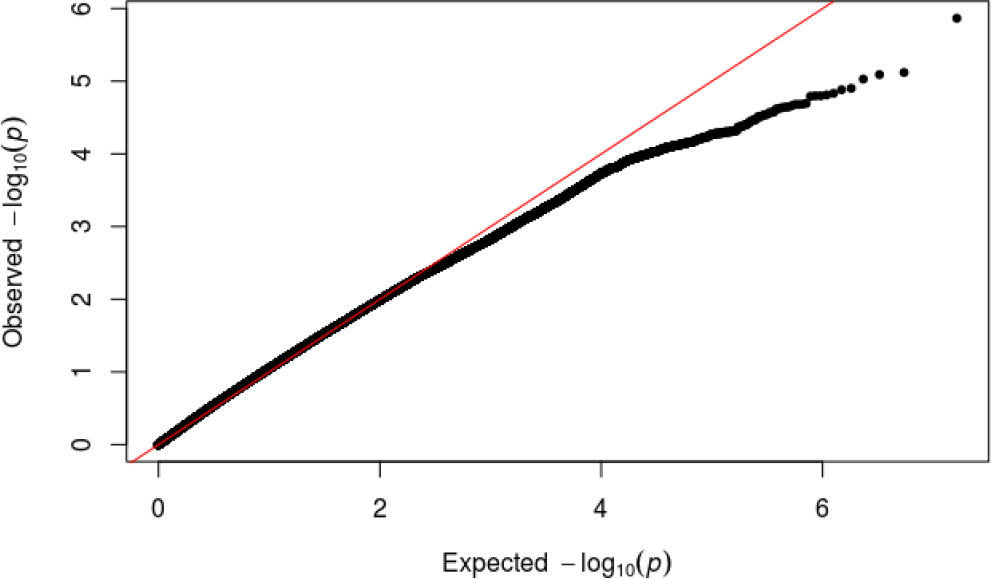
Observed versus expected distribution of p-values for the colorectal cancer genome-wide scan.

**Fig. 7.**
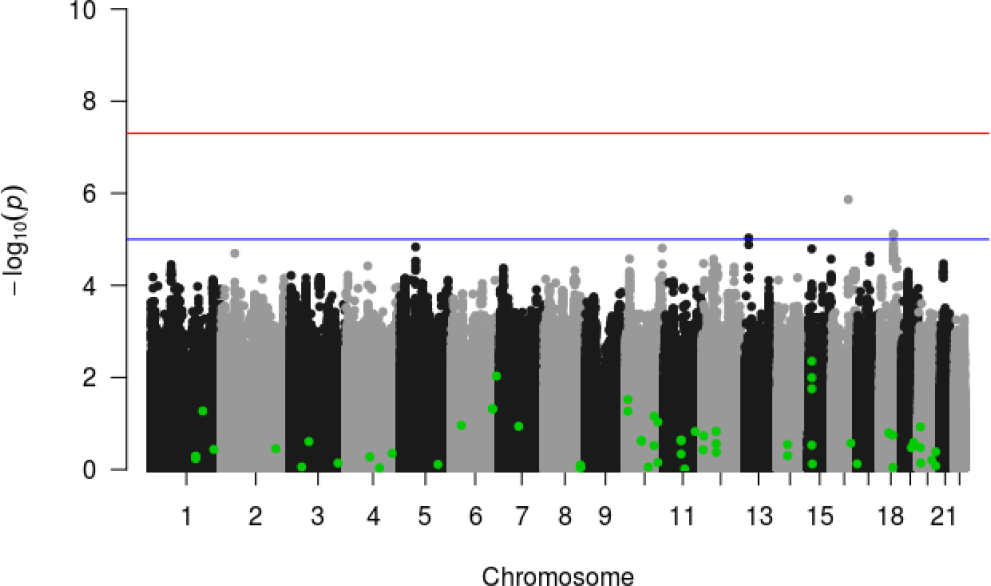
Manhattan plot for colorectal cancer genome-wide scan. Several SNPs were flagged with p-values < 1E-5. Green dots indicate variants flagged in previous genetic association studies.

**Table 4.** Results from colorectal cancer genome-wide scan. Genetic variants with a marginal p-value < 1E-5.

| Rsid | Gene | Chr | Pos | Ref | Alt | MaF | OR | SE | P |
| --- | --- | --- | --- | --- | --- | --- | --- | --- | --- |
| rs201447553 | Intergenic | 13 | 32081199 | G | GT | 0.218 | 10.746 | 0.536 | 9.36E-06 |
| - | Intergenic | 16 | 59740698 | T | TA | 0.423 | 17.539 | 0.593 | 1.36E-06 |
| rs17663205 | Utr3:MRO | 18 | 48324484 | G | C | 0.169 | 0.089 | 0.540 | 7.59E-06 |
| rs56387261 | Intron:MRO | 18 | 48325957 | C | T | 0.170 | 0.088 | 0.545 | 8.12E-06 |

Chr: Chromosome
Pos: Chromosome Position (build 37)
Ref/Alt: Reference and alternate allele
MaF: Minor allele frequency
OR: Odds ratio
SE: Standard error

## References

1. L. A. Torre, F. Bray, R. L. Siegel, J. Ferlay, J. Lortet-Tieulent, and A. Jemal, “Global cancer statistics, 2012,” CA: a cancer journal for clinicians, vol. 65, no. 2, pp. 87–108, 2015.

2. R. L. Siegel, K. D. Miller, and A. Jemal, “Cancer statistics, 2016,” CA: a cancer journal for clinicians, vol. 66, no. 1, pp. 7–30, 2016.

3. B. Pardamean, J. W. Baurley, C. I. Pardamean, and J. C. Figueiredo, “Changing col-orectal cancer trends in asians,” International journal of colorectal disease, vol. 31, no. 8, p. 1537, 2016.

4. M. A. Pourhoseingholi, “Increased burden of colorectal cancer in asia,” World journal of gastrointestinal oncology, vol. 4, no. 4, p. 68, 2012.

5. M. P. W. journal of gastrointestinal Oncology and undefined 2012, “Increased burden of colorectal cancer in Asia,” ncbi.nlm.nih.gov.

6. C. J. Ng, C. H. Teo, N. Abdullah, W. P. Tan, and H. M. Tan, “Relationships between cancer pattern, country income and geographical region in Asia.,” BMC cancer, vol. 15, p. 613, sep 2015.

7. Globocan, “Estimated Cancer Incidence, Mortality, and Prevalence Worldwide in 2012.”

8. U. Peters, S. Bien, and N. Zubair, “Genetic architecture of colorectal cancer,” Gut, vol. 64, no. 10, pp. 1623–1636, 2015.

9. C. A. Haiman and D. O. Stram, “Exploring genetic susceptibility to cancer in diverse populations,” Current opinion in genetics & development, vol. 20, no. 3, pp. 330–335, 2010.

10. W.-H. Jia, B. Zhang, K. Matsuo, A. Shin, Y.-B. Xiang, S. H. Jee, D.-H. Kim, Z. Ren, Q. Cai, J. Long, et al., “Genome-wide association analyses in east asians identify new susceptibility loci for colorectal cancer,” Nature genetics, vol. 45, no. 2, p. 191, 2013.

11. B. Zhang, W.-H. Jia, K. Matsuda, S.-S. Kweon, K. Matsuo, Y.-B. Xiang, A. Shin, S. H. Jee, D.-H. Kim, Q. Cai, et al., “Large-scale genetic study in east asians identifies six new loci associated with colorectal cancer risk,” Nature genetics, vol. 46, no. 6, p. 533, 2014.

12. J. W. Baurley, C. K. Edlund, C. I. Pardamean, D. V. Conti, and A. W. Bergen, “Smoke-screen: A targeted genotyping array for addiction research,” BMC Genomics, vol. 17, p. 145, dec 2016.

13. S. Das, L. Forer, S. Schönherr, C. Sidore, A. E. Locke, A. Kwong, S. I. Vrieze, E. Y. Chew, S. Levy, M. McGue, et al., “Next-generation genotype imputation service and methods,” Nature genetics, vol. 48, no. 10, p. 1284, 2016.

14. G. P. Consortium et al., “A global reference for human genetic variation,” Nature, vol. 526, no. 7571, p. 68, 2015.

15. P. Loh, “Eagle v2.4 user manual.”

16. A. Raj, M. Stephens, and J. K. Pritchard, “faststructure: variational inference of population structure in large snp data sets,” Genetics, vol. 197, no. 2, pp. 573–589, 2014.

17. S. L. Schmit, F. R. Schumacher, C. K. Edlund, D. V. Conti, U. Ihenacho, P. Wan, D. Van Den Berg, G. Casey, B. K. Fortini, H. J. Lenz, T. Tusié-Luna, C. A. Aguilar-Salinas, H. Moreno-Macías, A. Huerta-Chagoya, M. L. Ordóñez-Sánchez, R. Rodríguez-Guillén, I. Cruz-Bautista, M. Rodríguez-Torres, L. L. Munñóz-Hernández, O. Arellano-Campos, D. Gómez, U. Alvirde, C. Gonzólez-Villalpando, M. E. González-Villalpando, L. L. Marchand, C. A. Haiman, and J. C. Figueiredo, “Genome-wide association study of colorectal cancer in Hispanics,” Carcinogenesis, vol. 37, pp. 547–556, jun 2016.

18. A. E. Raftery, “Approximate bayes factors and accounting for model uncertainty in generalised linear models,” Biometrika, vol. 83, no. 2, pp. 251–266, 1996.

19. A. Armagan, D. B. Dunson, and J. Lee, “Generalized double pareto shrinkage,” Statistica Sinica, vol. 23, no. 1, p. 119, 2013.

20. A. P. Dempster, N. M. Laird, and D. B. Rubin, “Maximum likelihood from incomplete data via the em algorithm,” Journal of the royal statistical society. Series B (methodological), pp. 1–38, 1977.

21. J. Friedman, T. Hastie, and R. Tibshirani, “Regularization paths for generalized linear models via coordinate descent,” Journal of statistical software, vol. 33, no. 1, p. 1, 2010.

22. N. G. Polson and J. G. Scott, “Data augmentation for non-gaussian regression models using variance-mean mixtures,” Biometrika, vol. 100, no. 2, pp. 459–471, 2013.

23. S. Konishi and G. Kitagawa, “Bayesian information criteria,” Information Criteria and Statistical Modeling, pp. 211–237, 2008.

24. R Core Team, R: A Language and Environment for Statistical Computing. R Foundation for Statistical Computing, Vienna, Austria, 2016.

25. S. Widjaja and H. Yo, “RM-049Colorectal cancer in Indonesia - a centre report,” Annals of Oncology, vol. 27, no. suppl 2, pp. ii97.2–ii97, 2016.

26. A. I. Phipps, N. M. Lindor, M. A. Jenkins, J. A. Baron, A. K. Win, S. Gallinger, R. Gryfe, and P. A. Newcomb, “Colon and rectal cancer survival by tumor location and microsatellite instability: the Colon Cancer Family Registry.,” Diseases of the colon and rectum, vol. 56, pp. 937–44, aug 2013.

27. K. Hemminki, I. Santi, M. Weires, H. Thomsen, J. Sundquist, and J. L. Bermejo, “Tumor location and patient characteristics of colon and rectal adenocarcinomas in relation to survival and TNM classes,” BMC Cancer, vol. 10, p. 688, dec 2010.

28. R. Cui, Y. Okada, S. G. Jang, J. L. Ku, J. G. Park, Y. Kamatani, N. Hosono, T. Tsunoda, V. Kumar, C. Tanikawa, N. Kamatani, R. Yamada, M. Kubo, Y. Nakamura, and K. Matsuda, “Common variant in 6q26-q27 is associated with distal colon cancer in an asian population,” Gut, vol. 60, pp. 799–805, June 2011.

29. L. Zhu, M. Du, D. Gu, L. Ma, H. Chu, N. Tong, J. Chen, Z. Zhang, and M. Wang, “Genetic variant rs7758229 in 6q26-q27 is not associated with colorectal cancer risk in a chinese population,” PLoS One, vol. 8, p. e59256, Mar. 2013.

30. C.-M. Hsu, P.-M. Lin, J.-G. Chang, H.-C. Lin, S.-H. Li, S.-F. Lin, and M.-Y. Yang, “Upregulated SLC22A3 has a potential for improving survival of patients with head and neck squamous cell carcinoma receiving cisplatin treatment,” Oncotarget, vol. 8, pp. 74348–74358, Sept. 2017.

31. C. Grisanzio, L. Werner, D. Takeda, B. C. Awoyemi, M. M. Pomerantz, H. Yamada, P. Sooriakumaran, B. D. Robinson, R. Leung, A. C. Schinzel, I. Mills, H. Ross-Adams, D. E. Neal, M. Kido, T. Yamamoto, G. Petrozziello, E. C. Stack, R. Lis, P. W. Kantoff, M. Loda, O. Sartor, S. Egawa, A. K. Tewari, W. C. Hahn, and M. L. Freedman, “Genetic and functional analyses implicate the NUDT11, HNF1B, and SLC22A3 genes in prostate cancer pathogenesis,” Proc. Natl. Acad. Sci. U. S. A., vol. 109, pp. 11252–11257, July 2012.

32. Q. Li and Y. Shu, “Role of solute carriers in response to anticancer drugs,” Mol Cell Ther, vol. 2, p. 15, May 2014.

33. S. Yokoo, S. Masuda, A. Yonezawa, T. Terada, T. Katsura, and K.-I. Inui, “Significance of organic cation transporter 3 (SLC22A3) expression for the cytotoxic effect of oxaliplatin in colorectal cancer,” Drug Metab. Dispos., vol. 36, pp. 2299–2306, Nov. 2008.

34. N. Whiffin, F. J. Hosking, S. M. Farrington, C. Palles, S. E. Dobbins, L. Zgaga, A. Lloyd, B. Kinnersley, M. Gorman, A. Tenesa, P. Broderick, Y. Wang, E. Barclay, C. Hayward, L. Martin, D. D. Buchanan, A. K. Win, J. Hopper, M. Jenkins, N. M. Lindor, P. A. Newcomb, S. Gallinger, D. Conti, F. Schumacher, G. Casey, T. Liu, Swedish Low-Risk Colorectal Cancer Study Group, H. Campbell, A. Lindblom, R. S. Houlston, I. P. Tomlinson, and M. G. Dunlop, “Identification of susceptibility loci for colorectal cancer in a genome-wide meta-analysis,” Hum. Mol. Genet., vol. 23, pp. 4729–4737, Sept. 2014.

35. C. Tanikawa, Y. Kamatani, A. Takahashi, Y. Momozawa, K. Leveque, S. Nagayama, K. Mimori, M. Mori, H. Ishii, J. Inazawa, J. Yasuda, A. Tsuboi, A. Shimizu, M. Sasaki, T. Yamaji, N. Sawada, M. Iwasaki, S. Tsugane, M. Naito, K. Wakai, T. Koyama, T. Takezaki, K. Yuji, Y. Murakami, Y. Nakamura, M. Kubo, and K. Matsuda, “GWAS identifies two novel colorectal cancer loci at 16q24.1 and 20q13.12,” Carcinogenesis, vol. 39, pp. 652–660, May 2018.

36. S. L. Schmit, C. K. Edlund, F. R. Schumacher, J. Gong, T. A. Harrison, J. R. Huyghe, C. Qu, M. Melas, D. J. Van Den Berg, H. Wang, S. Tring, S. J. Plummer, D. Albanes, M. H. Alonso, C. I. Amos, K. Anton, A. K. Aragaki, V. Arndt, E. L. Barry, S. I. Berndt, S. Bezieau, S. Bien, A. Bloomer, J. Boehm, M.-C. Boutron-Ruault, H. Brenner, S. Brezina, D. D. Buchanan, K. Butterbach, B. J. Caan, P. T. Campbell, C. S. Carlson, J. E. Castelao, A. T. Chan, J. Chang-Claude, S. J. Chanock, I. Cheng, Y.-W. Cheng, L. S. Chin, J. M. Church, T. Church, G. A. Coetzee, M. Cotterchio, M. Cruz Correa, K. R. Curtis, D. Duggan, D. F. Easton, D. English, E. J. M. Feskens, R. Fischer, L. M. FitzGerald, B. K. Fortini, L. G. Fritsche, C. S. Fuchs, M. Gago-Dominguez, M. Gala, S. J. Gallinger, W. J. Gauderman, G. G. Giles, E. L. Giovannucci, S. M. Gogarten, C. Gonzalez-Villalpando, E. M. Gonzalez-Villalpando, W. M. Grady, J. K. Greenson, A. Gsur, M. Gunter, C. A. Haiman, J. Hampe, S. Harlid, J. F. Harju, R. B. Hayes, P. Hofer, M. Hoffmeister, J. L. Hopper, S.-C. Huang, J. M. Huerta, T. J. Hudson, D. J. Hunter, G. E. Idos, M. Iwasaki, R. D. Jackson, E. J. Jacobs, S. H. Jee, M. A. Jenkins, W.-H. Jia, S. Jiao, A. D. Joshi, L. N. Kolonel, S. Kono, C. Kooperberg, V. Krogh, T. Kuehn, S. Küry, A. LaCroix, C. A. Laurie, F. Lejbkowicz, M. Lemire, H.-J. Lenz, D. Levine, C. I. Li, L. Li, W. Lieb, Y. Lin, N. M. Lindor, Y.-R. Liu, F. Loupakis, Y. Lu, F. Luh, J. Ma, C. Mancao, F. J. Manion, S. D. Markowitz, V. Martin, K. Matsuda, K. Matsuo, K. J. McDonnell, C. E. McNeil, R. Milne, A. J. Molina, B. Mukherjee, N. Murphy, P. A. Newcomb, K. Offit, H. Omichessan, D. Palli, J. P. P. Cotoré, J. Pérez-Mayoral, P. D. Pharoah, J. D. Potter, C. Qu, L. Raskin, G. Rennert, H. S. Rennert, B. M. Riggs, C. Schafmayer, R. E. Schoen, T. A. Sellers, D. Seminara, G. Severi, W. Shi, D. Shibata, X.-O. Shu, E. M. Siegel, M. L. Slattery, M. Southey, Z. K. Stadler, M. C. Stern, S. Stintzing, D. Taverna, S. N. Thibodeau, D. C. Thomas, A. Trichopoulou, S. Tsugane, C. M. Ulrich, F. J. B. van Duijnhoven, B. van Guelpan, J. Vijai, J. Virtamo, S. J. Weinstein, E. White, A. K. Win, A. Wolk, M. Woods, A. H. Wu, K. Wu, Y.-B. Xiang, Y. Yen, B. W. Zanke, Y.-X. Zeng, B. Zhang, N. Zubair, S.-S. Kweon, J. C. Figueiredo, W. Zheng, L. L. Marchand, A. Lindblom, V. Moreno, U. Peters, G. Casey, L. Hsu, D. V. Conti, and S. B. Gruber, “Novel common genetic susceptibility loci for colorectal cancer,” J. Natl. Cancer Inst., June 2018.

37. F. R. Schumacher, S. L. Schmit, S. Jiao, C. K. Edlund, H. Wang, B. Zhang, L. Hsu, S.-C. Huang, C. P. Fischer, J. F. Harju, G. E. Idos, F. Lejbkowicz, F. J. Manion, K. McDonnell, C. E. McNeil, M. Melas, H. S. Rennert, W. Shi, D. C. Thomas, D. J. Van Den Berg, C. M. Hutter, A. K. Aragaki, K. Butterbach, B. J. Caan, C. S. Carlson, S. J. Chanock, K. R. Curtis, C. S. Fuchs, M. Gala, E. L. Giovannucc, S. M. Gogarten, R. B. Hayes, B. Henderson, D. J. Hunter, R. D. Jackson, L. N. Kolonel, C. Kooperberg, Küry, A. LaCroix, C. C. Laurie, C. A. Laurie, M. Lemire, D. Levine, J. Ma, K. W. Makar, C. Qu, D. Taverna, C. M. Ulrich, K. Wu, S. Kono, D. W. West, S. I. Berndt, S. Bezieau, H. Brenner, P. T. Campbell, A. T. Chan, J. Chang-Claude, G. A. Coetzee, D. V. Conti, D. Duggan, J. C. Figueiredo, B. K. Fortini, S. J. Gallinger, W. J. Gauderman, G. Giles, R. Green, R. Haile, T. A. Harrison, M. Hoffmeister, J. L. Hopper, T. J. Hudson, E. Jacobs, M. Iwasaki, S. H. Jee, M. Jenkins, W.-H. Jia, A. Joshi, L. Li, N. M. Lindor, K. Matsuo, V. Moreno, B. Mukherjee, P. A. Newcomb, J. D. Potter, L. Raskin, G. Rennert, S. Rosse, G. Severi, R. E. Schoen, D. Seminara, X.-O. Shu, M. L. Slat-tery, S. Tsugane, E. White, Y.-B. Xiang, B. W. Zanke, W. Zheng, L. Le Marchand, Casey, S. B. Gruber, and U. Peters, “Genome-wide association study of colorectal cancer identifies six new susceptibility loci,” Nat. Commun., vol. 6, p. 7138, July 2015.

38. J. B. Sneddon, H. H. Zhen, K. Montgomery, M. van de Rijn, A. D. Tward, R. West, Gladstone, H. Y. Chang, G. S. Morganroth, A. E. Oro, and P. O. Brown, “Bone morphogenetic protein antagonist gremlin 1 is widely expressed by cancer-associated stromal cells and can promote tumor cell proliferation,” Proc. Natl. Acad. Sci. U. S. A., vol. 103, pp. 14842–14847, Oct. 2006.

39. H. Stabile, S. Mitola, E. Moroni, M. Belleri, S. Nicoli, D. Coltrini, F. Peri, A. Pessi, L. Orsatti, F. Talamo, V. Castronovo, D. Waltregny, F. Cotelli, D. Ribatti, and Presta, “Bone morphogenic protein antagonist drm/gremlin is a novel proangiogenic factor,” Blood, vol. 109, pp. 1834–1840, Mar. 2007.

40. J. Ziai, E. Matloff, J. Choi, N. Kombo, M. Materin, and A. E. Bale, “Defining the polyposis/colorectal cancer phenotype associated with the ashkenazi GREM1 duplication: counselling and management recommendations,” Genet. Res., vol. 98, p. e5, Mar. 2016.

41. H. Davis, S. Irshad, M. Bansal, H. Rafferty, T. Boitsova, C. Bardella, E. Jaeger, A. Lewis, L. Freeman-Mills, F. C. Giner, P. Rodenas-Cuadrado, S. Mallappa, S. Clark, H. Thomas, R. Jeffery, R. Poulsom, M. Rodriguez-Justo, M. Novelli, R. Chetty, A. Silver, O. J. Sansom, F. R. Greten, L. M. Wang, J. E. East, I. Tomlinson, and S. J. Leedham, “Aberrant epithelial GREM1 expression initiates colonic tumorigenesis from cells outside the stem cell niche,” Nat. Med., vol. 21, pp. 62–70, Jan. 2015.

42. F. O. Desmet, D. Hamroun, M. Lalande, G. Collod-Bëroud, M. Claustres, and C. Béroud, “Human Splicing Finder: An online bioinformatics tool to predict splicing signals,” Nucleic Acids Research, vol. 37, no. 9, 2009.

43. Y. Yang, P. Junjie, C. Sanjun, and Y. Ma, “Long non-coding RNAs in Colorectal Cancer: Progression and Future Directions,” Journal of Cancer, vol. 8, 2017.

44. W. Guo, Q. Wang, Y. Zhan, X. Chen, Q. Yu, J. Zhang, Y. Wang, X. J. Xu, and L. Zhu, “Transcriptome sequencing uncovers a three-long noncoding RNA signature in predicting breast cancer survival,” Scientific Reports, vol. 6, 2016.

45. E. S. Wan, W. Qiu, V. J. Carey, J. Morrow, H. Bacherman, M. G. Foreman, J. E. Hokanson, R. P. Bowler, J. D. Crapo, and D. L. DeMeo, “Smoking-associated sitespecific differential methylation in buccal mucosa in the COPDGene study,” American Journal of Respiratory Cell and Molecular Biology, vol. 53, pp. 246–254, aug 2015.

46. T.-H. Liu, F. Zheng, M.-Y. Cai, L. Guo, H.-X. Lin, J.-W. Chen, Y.-J. Liao, H.-F. Kung, Y.-X. Zeng, and D. Xie, “The putative tumor activator ARHGEF3 promotes nasopharyngeal carcinoma cell pathogenesis by inhibiting cellular apoptosis,” Oncotarget, vol. 7, pp. 25836–25848, May 2016.

47. Y. Shi, G.-B. Chen, X.-X. Huang, C.-X. Xiao, H.-H. Wang, Y.-S. Li, J.-F. Zhang, S. Li, Y. Xia, J.-L. Ren, and B. Guleng, “Dragon (repulsive guidance molecule b, RGMb) is a novel gene that promotes colorectal cancer growth,” Oncotarget, vol. 6, pp. 20540–20554, Aug. 2015.

48. I. Suryapranata and R. Kusuma unpublished, N.D.

49. U. Peters, S. Jiao, F. R. Schumacher, C. M. Hutter, A. K. Aragaki, J. A. Baron, S. I. Berndt, S. Bézieau, H. Brenner, K. Butterbach, B. J. Caan, P. T. Campbell, C. S. Carlson, G. Casey, A. T. Chan, J. Chang-Claude, S. J. Chanock, L. S. Chen, G. A. Coetzee, S. G. Coetzee, D. V. Conti, K. R. Curtis, D. Duggan, T. Edwards, C. S. Fuchs, S. Gallinger, E. L. Giovannucci, S. M. Gogarten, S. B. Gruber, R. W. Haile, T. A. Harrison, R. B. Hayes, B. E. Henderson, M. Hoffmeister, J. L. Hopper, T. J. Hudson, D. J. Hunter, R. D. Jackson, S. H. Jee, M. A. Jenkins, W. H. Jia, L. N. Kolonel, C. Kooperberg, S. Küry, A. Z. Lacroix, C. C. Laurie, C. A. Laurie, L. Le Marchand, M. Lemire, D. Levine, N. M. Lindor, Y. Liu, J. Ma, K. W. Makar, K. Matsuo, P. A. Newcomb, J. D. Potter, R. L. Prentice, C. Qu, T. Rohan, S. A. Rosse, R. E. Schoen, D. Seminara, M. Shrubsole, X. O. Shu, M. L. Slattery, D. Taverna, S. N. Thibodeau, C. M. Ulrich, E. White, Y. Xiang, B. W. Zanke, Y. X. Zeng, B. Zhang, W. Zheng, and L. Hsu, “Identification of genetic susceptibility loci for colorectal tumors in a genomewide meta-analysis,” Gastroenterology, vol. 144, pp. 799–807.e24, apr 2013.

50. N. Whiffin, F. J. Hosking, S. M. Farrington, C. Palles, S. E. Dobbins, L. Zgaga, A. Lloyd, B. Kinnersley, M. Gorman, A. Tenesa, P. Broderick, Y. Wang, E. Barclay, C. Hayward, L. Martin, D. D. Buchanan, A. K. Win, J. Hopper, M. Jenkins, N. M. Lindor, P. A. Newcomb, S. Gallinger, D. Conti, F. Schumacher, G. Casey, T. Liu, H. Campbell, A. Lindblom, R. S. Houlston, I. P. Tomlinson, and M. G. Dunlop, “Identification of susceptibility loci for colorectal cancer in a genome-wide meta-analysis,” Human Molecular Genetics, vol. 23, pp. 4729–4737, sep 2014.

51. R. S. Houlston, J. Cheadle, S. E. Dobbins, A. Tenesa, A. M. Jones, K. Howarth, S. L. Spain, P. Broderick, E. Domingo, S. Farrington, J. G. Prendergast, A. M. Pittman, E. Theodoratou, C. G. Smith, B. Olver, A. Walther, R. A. Barnetson, M. Churchman, E. E. Jaeger, S. Penegar, E. Barclay, L. Martin, M. Gorman, R. Mager, E. Johnstone, R. Midgley, I. Niittymäki, S. Tuupanen, J. Colley, S. Idziaszczyk, H. J. Thomas, A. M. Lucassen, D. G. R. Evans, E. R. Maher, T. Maughan, A. Dimas, E. Dermitzakis, J. B. Cazier, L. A. Aaltonen, P. Pharoah, D. J. Kerr, L. G. Carvajal-Carmona, H. Campbell, M. G. Dunlop, and I. P. Tomlinson, “Meta-analysis of three genome-wide association studies identifies susceptibility loci for colorectal cancer at 1q41, 3q26.2, 12q13.13 and 20q13.33,” Nature Genetics, vol. 42, pp. 973–977, nov 2010.

52. F. R. Schumacher, S. L. Schmit, S. Jiao, C. K. Edlund, H. Wang, B. Zhang, L. Hsu, S. C. Huang, C. P. Fischer, J. F. Harju, G. E. Idos, F. Lejbkowicz, F. J. Manion, K. McDonnell, C. E. McNeil, M. Melas, H. S. Rennert, W. Shi, D. C. Thomas, D. J. Van Den Berg, C. M. Hutter, A. K. Aragaki, K. Butterbach, B. J. Caan, C. S. Carlson, S. J. Chanock, K. R. Curtis, C. S. Fuchs, M. Gala, E. L. Giocannucci, S. M. Gogarten, R. B. Hayes, B. Henderson, D. J. Hunter, R. D. Jackson, L. N. Kolonel, C. Kooperberg, Kury, A. Lacroix, C. C. Laurie, C. A. Laurie, M. Lemire, D. Levine, J. Ma, K. W. Makar, C. Qu, D. Taverna, C. M. Ulrich, K. Wu, S. Kono, D. W. West, S. I. Berndt, S. Bezieau, H. Brenner, P. T. Campbell, A. T. Chan, J. Chang-Claude, G. A. Coetzee, D. V. Conti, D. Duggan, J. C. Figueiredo, B. K. Fortini, S. J. Gallinger, W. J. Gauderman, G. Giles, R. Green, R. Haile, T. A. Harrison, M. Hoffmeister, J. L. Hopper, T. J. Hudson, E. Jacobs, M. Iwasaki, S. H. Jee, M. Jenkins, W. H. Jia, A. Joshi, L. Li, N. M. Lindor, K. Matsuo, V. Moreno, B. Mukherjee, P. A. Newcomb, J. D. Potter, L. Raskin, G. Rennert, S. Rosse, G. Severi, R. E. Schoen, D. Seminara, X. O. Shu, M. L. Slattery, S. Tsugane, E. White, Y. B. Xiang, B. W. Zanke, W. Zheng, L. Le Marchand, G. Casey, S. B. Gruber, and U. Peters, “Genome-wide association study of colorectal cancer identifies six new susceptibility loci,” Nature Communications, vol. 6, p. 7138, dec 2015.

53. L. M. Real, A. Ruiz, J. Gayán, A. González-Pérez, M. E. Sáez, R. Ramírez-Lorca, F. J. Morón, J. Velasco, R. Marginet-Flinch, E. Musulén, J. M. Carrasco, C. Moreno-Rey, E. Vázquez, M. Chaves-Conde, J. A. Moreno-Nogueira, M. Hidalgo-Pascual, E. Ferrero-Herrero, S. Castellví-Bel, A. Castells, C. Fernandez-Rozadilla, C. Ruiz-Ponte, A. Carracedo, B. González, S. Alonso, and M. Perucho, “A colorectal cancer susceptibility new variant at 4q26 in the Spanish population identified by genome-wide association analysis,” PLoS ONE, vol. 9, p. e101178, jun 2014.

54. M. G. Dunlop, S. E. Dobbins, S. M. Farrington, A. M. Jones, C. Palles, N. Whiffin, A. Tenesa, S. Spain, P. Broderick, L. Y. Ooi, E. Domingo, C. Smillie, M. Henrion, M. Frampton, L. Martin, G. Grimes, M. Gorman, C. Semple, Y. P. Ma, E. Barclay, J. Prendergast, J. B. Cazier, B. Olver, S. Penegar, S. Lubbe, I. Chander, L. G. Carvajal-Carmona, S. Ballereau, A. Lloyd, J. Vijayakrishnan, L. Zgaga, I. Rudan, E. Theodoratou, H. Thomas, E. Maher, G. Evans, L. Walker, D. Halliday, A. Lucassen, J. Paterson, S. Hodgson, T. Homfray, L. Side, L. Izatt, A. Donaldson, S. Tomkins, P. Morrison, C. Brewer, A. Henderson, R. Davidson, V. Murday, J. Cook, N. Haites, T. Bishop, E. Sheridan, A. Green, C. Marks, S. Carpenter, M. Broughton, L. Greenhalge, M. Suri, J. M. Starr, I. Deary, I. Kirac, D. Kovacevia, L. A. Aaltonen, L. Renkonen-Sinisalo, J. P. Mecklin, K. Matsuda, Y. Nakamura, Y. Okada, S. Gallinger, D. J. Duggan, D. Conti, P. Newcomb, J. Hopper, M. A. Jenkins, F. Schumacher, G. Casey, D. Easton, M. Shah, P. Pharoah, A. Lindblom, T. Liu, D. Edler, C. Lenander, J. Dalén, F. Hjern, N. Lundqvist, U. Lindforss, L. Påhlman, K. Smedh, A. Törnqvist, J. Holm, M. Janson, M. Andersson, S. Ekelund, L. Olsson, C. G. Smith, H. West, J. P. Cheadle, G. MacDonald, L. M. Samuel, A. Ahmad, P. Corrie, D. Jodrell, C. Palmer, C. Wilson, J. O’Hagan, D. Smith, R. McDermott, J. Walshe, J. Cassidy, A. McDonald, N. Mohammed, J. White, H. Yosef, O. Breathnach, L. Grogan, R. Thomas, M. Eatock, P. Henry, R. Houston, P. Johnston, R. Wilson, I. Geh, F. Danwata, A. Hindley, S. Susnerwala, C. Bradley, A. Conn, A. Raine, C. Twelves, S. Falk, K. Hopkins, S. Tahir, A. Dhadda, A. Maraveyas, J. Sgouros, M. Teo, R. Ahmad, S. Cleator, A. Creak, C. Lowdell, P. Riddle, K. Benstead, D. Farrugia, N. Reed, S. Shepherd, E. Levine, S. Mullamitha, M. Saunders, J. Valle, G. Wilson, A. Jones, A. Weaver, P. I. Clark, B. Haylock, M. I. Iqbal, A. S. Myint, S. Beesley, T. Sevitt, J. Nicoll, F. Daniel, V. Ford, T. Talbot, M. Butt, A. Hamid, P. MacK, R. Roy, R. Osborne, F. McKinna, H. Alsab, D. Basu, P. Murray, B. Sizer, F. A. Azam, R. Neupane, A. Waterston, J. Glaholm, C. Blesing, S. Lowndes, A. Medisetti, A. Gaya, M. Leslie, N. Maisey, P. Ross, G. Dunn, O. Al-Salihi, H. S. Wasan, L. T. Tan, J. Dent, U. Hofmann, J. K. Joffe, E. Sherwin, R. Soomal, A. Chakrabarti, S. Joseph, J. Van Der Voet, N. J. Wadd, D. Wilson, S. Anjarwalia, J. Hall, R. Hughes, A. Polychronis, J. H. Scarffe, M. Hill, R. D. James, R. Shah, J. Summers, A. Hartley, D. Carney, J. McCaffrey, B. Bystricky, S. O’Reilly, R. Gupta, T. Al-Mishlab, F. Gidden, R. O’Hara, J. Stewart, R. Ashford, R. Glynne-Jones, M. Harrison, S. Mawdsley, H. Barlow, M. Tighe, J. Walther, J. Neal, C. Rees, J. Bridgewater, S. Karp, U. McGovern, P. J. Atherton, H. El-Deeb, C. MacMillan, K. Patel, E. M. Bessell, P. D. Dickinson, V. Potter, C. Jephcott, K. McAdam, J. Wrigley, S. Muthuramalingam, A. O’Callaghan, L. Melcher, C. Braconi, J. I. Geh, D. Palmer, P. Narayana, N. Steven, A. Gaya, S. Rudman, P. Chakraborti, K. Kelly, C. MacGregor, D. Whillis, A. Freebairn, J. Gilder-sleve, S. Sharif, G. Astras, T. Hickish, D. Beech, R. Ellis, R. Kulkarni, K. Shankland, R. Begent, A. Mayer, T. Meyer, S. Strauss, V. Hall, S. Raj, I. Chau, D. Cunningham, A. Birtle, A. Biswas, M. Wise, S. Cummins, S. Essapen, G. Middleton, C. Topham, R. Langley, A. Webb, M. Wilkins, T. J. Iveson, C. Askill, J. Wagstaff, A. Azzabi, A. Bateman, J. Prejbisz, D. Tsang, N. Ali, A. Jones, P. O’Neill, C. Cottrill, D. Propper, F. J. Lofts, J. Kennedy, D. A. Anthoney, R. Cooper, A. Crellin, A. Melcher, M. Seymour, C. Baughan, E. Alexander, J. Crown, D. Fennelly, F. Adab, S. Giridharan, I. Pedley, K. Wright, P. Bliss, G. Cogill, N. Lo, E. Toy, D. Hochhauser, J. Ledermann, A. Brewster, T. Maughan, D. Mort, S. Mukherjee, W. Dobrowsky, P. Calvert, G. Leonard, H. Ford, A. M. Moody, S. Goriah, M. Wilkins, S. Clive, L. Dawson, C. McLean, H. A. Phillips, K. Gopi, M. Tomlinson, S. Clenton, D. Furniss, J. Hornbuckle, S. Pledge, J. Wadsley, M. Abbas, E. Marshall, C. Harper-Wynne, A. Barnes, S. Kumar, V. Vigneswaran, S. Gollins, M. Genton, G. Sparrow, C. Bale, C. Fuller, A. Mullard, N. Stuart, R. Williams, M. Keane, T. Maughen, R. Adams, A. Madi, E. Hodgkinson, P. Rogers, M. Pope, R. Kaplan, A. Meade, M. Parmar, S. Kenny, D. Fisher, L. Harper, J. Mitchell, L. Nichols, B. Sydes, L. Clement, E. Kay, C. Courtney, M. Gallagher, C. Murphy, L. Thompson, S. Beall, S. Hassan, R. Gracie, G. Griffiths, M. Mason, C. Parker, R. Rudd, P. Johnson, J. Whelan, J. Northover, J. Brown, M. Aapro, R. Stout, R. Midgley, D. J. Kerr, H. Campbell, I. P. Tomlinson, and R. S. Houlston, “Common variation near CDKN1A, POLD3 and SHROOM2 influences colorectal cancer risk,” Nature Genetics, vol. 44, pp. 770–776, jul 2012.

55. R. Cui, Y. Okada, S. G. Jang, J. L. Ku, J. G. Park, Y. Kamatani, N. Hosono, T. Tsunoda, V. Kumar, C. Tanikawa, N. Kamatani, R. Yamada, M. Kubo, Y. Nakamura, and K. Matsuda, “Common variant in 6q26-q27 is associated with distal colon cancer in an Asian population,” Gut, vol. 60, pp. 799–805, jun 2011.

56. B. W. Zanke, C. M. Greenwood, J. Rangrej, R. Kustra, A. Tenesa, S. M. Farrington, J. Prendergast, S. Olschwang, T. Chiang, E. Crowdy, V. Ferretti, P. Laflamme, S. Sundararajan, S. Roumy, J. F. Olivier, F. Robidoux, R. Sladek, A. Montpetit, P. Campbell, S. Bezieau, A. M. O’Shea, G. Zogopoulos, M. Cotterchio, P. Newcomb, J. McLaughlin, B. Younghusband, R. Green, J. Green, M. E. Porteous, H. Campbell, H. Blanche, M. Sahbatou, E. Tubacher, C. Bonaiti-Pellié, B. Buecher, E. Riboli, S. Kury, S. J. Chanock, J. Potter, G. Thomas, S. Gallinger, T. J. Hudson, and M. G. Dunlop, “Genome-wide association scan identifies a colorectal cancer susceptibility locus on chromosome 8q24,” Nature Genetics, vol. 39, pp. 989–994, aug 2007.

57. S. B. Gruber, V. Moreno, L. S. Rozek, H. S. Rennerts, F. Lejbkowicz, J. D. Bonner, J. K. Greenson, T. J. Giordano, E. R. Fearson, and G. Rennert, “Genetic variation in 8q24 associated with risk of colorectal cancer.,” Cancer biology & therapy, vol. 6, pp. 1143–7, jul 2007.

58. C. A. Haiman, L. Le Marchand, J. Yamamato, D. O. Stram, X. Sheng, L. N. Kolonel, A. H. Wu, D. Reich, and B. E. Henderson, “A common genetic risk factor for colorectal and prostate cancer,” Nature Genetics, vol. 39, pp. 954–956, aug 2007.

59. I. P. Tomlinson, E. Webb, L. Carvajal-Carmona, P. Broderick, K. Howarth, A. M. Pittman, S. Spain, S. Lubbe, A. Walther, K. Sullivan, E. Jaeger, S. Fielding, A. Rowan, J. Vijayakrishnan, E. Domingo, I. Chandler, Z. Kemp, M. Qureshi, S. M. Farrington, A. Tenesa, J. G. Prendergast, R. A. Barnetson, S. Penegar, E. Barclay, W. Wood, L. Martin, M. Gorman, H. Thomas, J. Peto, D. T. Bishop, R. Gray, E. R. Maher, A. Lucassen, D. Kerr, D. G. R. Evans, C. Schafmayer, S. Buch, H. Völzke, J. Hampe, S. Schreiber, U. John, T. Koessler, P. Pharoah, T. Van Wezel, H. Morreau, J. T. Wijnen, J. L. Hopper, M. C. Southey, G. G. Giles, G. Severi, S. Castellví-Bel, C. Ruiz-Ponte, A. Carracedo, A. Castells, A. Försti, K. Hemminki, P. Vodicka, A. Naccarati, L. Lipton, J. W. Ho, K. K. Cheng, P. C. Sham, J. Luk, J. A. Agúndez, J. M. Ladero, M. De La Hoya, T. Caldés, I. Niittymäki, S. Tuupanen, A. Karhu, L. Aaltonen, J. B. Cazier, H. Campbell, M. G. Dunlop, and R. S. Houlston, “A genome-wide association study identifies colorectal cancer susceptibility loci on chromosomes 10p14 and 8q23.3,” Nature Genetics, vol. 40, pp. 623–630, may 2008.

60. C. M. Hutter, M. L. Slattery, D. J. Duggan, J. Muehling, K. Curtin, L. Hsu, S. A. Beresford, A. Rajkovic, G. E. Sarto, J. R. Marshall, N. Hammad, R. Wallace, K. W. Makar, R. L. Prentice, B. J. Caan, J. D. Potter, and U. Peters, “Characterization of the association between 8q24 and colon cancer: Gene-environment exploration and metaanalysis,” BMC Cancer, vol. 10, p. 670, dec 2010.

61. A. Tenesa, S. M. Farrington, J. G. Prendergast, M. E. Porteous, M. Walker, N. Haq, R. A. Barnetson, E. Theodoratou, R. Cetnarskyj, N. Cartwright, C. Semple, A. J. Clark, F. J. Reid, L. A. Smith, K. Kavoussanakis, T. Koessler, P. D. Pharoah, S. Buch, C. Schafmayer, J. Tepel, S. Schreiber, H. Völzke, C. O. Schmidt, J. Hampe, J. Chang-Claude, M. Hoffmeister, H. Brenner, S. Wilkening, F. Canzian, G. Capella, V. Moreno, I. J. Deary, J. M. Starr, I. P. Tomlinson, Z. Kemp, K. Howarth, L. Carvajal-Carmona, E. Webb, P. Broderick, J. Vijayakrishnan, R. S. Houlston, G. Rennert, D. Ballinger, L. Rozek, S. B. Gruber, K. Matsuda, T. Kidokoro, Y. Nakamura, B. W. Zanke, C. M. Greenwood, J. Rangrej, R. Kustra, A. Montpetit, T. J. Hudson, S. Gallinger, H. Campbell, and M. G. Dunlop, “Genome-wide association scan identifies a colorectal cancer susceptibility locus on 11q23 and replicates risk loci at 8q24 and 18q21,” Nature Genetics, vol. 40, pp. 631–637, may 2008.

62. H. Wang, C. A. Haiman, T. Burnett, B. K. Fortini, L. N. Kolonel, B. E. Henderson, L. B. Signorello, W. J. Blot, T. O. Keku, S. I. Berndt, P. A. Newcomb, M. Pande, C. I. Amos, D. W. West, G. Casey, R. S. Sandler, R. Haile, D. O. Stram, and L. Le Marchand, “Fine-mapping of genome-wide association study-identified risk loci for colorectal cancer in African Americans,” Human Molecular Genetics, vol. 22, pp. 5048–5055, dec 2013.

63. E. Jaeger, E. Webb, K. Howarth, L. Carvajal-Carmona, A. Rowan, P. Broderick, A. Walther, S. Spain, A. Pittman, Z. Kemp, K. Sullivan, K. Heinimann, S. Lubbe, E. Domingo, E. Barclay, L. Martin, M. Gorman, I. Chandler, J. Vijayakrishnan, W. Wood, E. Papaemmanuil, S. Penegar, M. Qureshi, S. Farrington, A. Tenesa, J. B. Cazier, D. Kerr, R. Gray, J. Peto, M. Dunlop, H. Campbell, H. Thomas, R. Houlston, and I. Tomlinson, “Common genetic variants at the CRAC1 (HMPS) locus on chromosome 15q13.3 influence colorectal cancer risk,” Nature Genetics, vol. 40, pp. 26–28, jan 2008.

64. P. Broderick, L. Carvajal-Carmona, A. M. Pittman, E. Webb, K. Howarth, A. Rowan, S. Lubbe, S. Spain, K. Sullivan, S. Fielding, E. Jaeger, J. Vijayakrishnan, Z. Kemp, M. Gorman, I. Chandler, E. Papaemmanuil, S. Penegar, W. Wood, G. Sellick, M. Qureshi, A. Teixeira, E. Domingo, E. Barclay, L. Martin, O. Sieber, D. Kerr, R. Gray, J. Peto, J. B. Cazier, I. Tomlinson, and R. S. Houlston, “A genome-wide association study shows that common alleles of SMAD7 influence colorectal cancer risk,” Nature Genetics, vol. 39, pp. 1315–1317, nov 2007.

